# Patient-derived gene and protein expression signatures of NGLY1 deficiency

**DOI:** 10.1101/2021.07.28.453930

**Authors:** Benedikt Rauscher, William F. Mueller, Sandra Clauder-Münster, Petra Jakob, M. Saiful Islam, Han Sun, Sonja Ghidelli-Disse, Markus Boesche, Marcus Bantscheff, Hannah Pflaumer, Paul Collier, Bettina Haase, Songjie Chen, Guangwen Wang, Vladimir Benes, Michael Snyder, Gerard Drewes, Lars M. Steinmetz

## Abstract

N-Glycanase 1 (NGLY1) deficiency is a rare and complex genetic disorder. Although recent studies have shed light on the molecular underpinnings of NGLY1 deficiency, a systematic characterization of gene and protein expression changes in patient-derived cells has been lacking. Here, we performed RNA-sequencing and mass spectrometry to determine the transcriptomes and proteomes of 66 cell lines representing 4 different cell types derived from 14 NGLY1 deficient patients and 17 controls. While gene and protein expression levels agreed well with each other, expression differences were more pronounced at the protein level. Although NGLY1 protein levels were up to 9.5-fold downregulated in patients compared to parent controls, depending on the genotype, NGLY1 protein was still detectable in all patient-derived lymphoblastoid cell lines. Consistent with the role of NGLY1 as a regulator of the transcription factor Nrf1, we observed a cell type-independent downregulation of proteasomal genes in NGLY1 deficient cells. In contrast, genes involved in ribosomal mRNA processing were upregulated in multiple cell types. In addition, we observed cell type-specific effects. For example, genes and proteins involved in glutathione synthesis, such as the glutamate-cystein ligase subunits GCLC and GCLM, were downregulated specifically in lymphoblastoid cells. We provide a web application that enables access to all results generated in this study at https://apps.embl.de/ngly1browser. This resource will guide future studies of NGLY1 deficiency in directions that are most relevant to patients.

## INTRODUCTION

N-glycanase (NGLY1) deficiency is a rare genetic disorder where both alleles of the NGLY1 gene are affected by deleterious mutations leading to a loss of function. Patients described with NGLY1 deficiency display a range of multi-organ symptoms including developmental delay, hypo-or alacrima, movement disorders, and seizures (Caglayan et al., 2015; Enns et al., 2014; Heeley and Shinawi, 2015; van Keulen et al., 2019; Lam et al., 2017; Need et al., 2012).

NGLY1 is a highly conserved protein that functions as part of the endoplasmic reticulum associated degradation pathway (ERAD) where it catalyzes the deglycosylation of misfolded N-glycoproteins before their degradation by the proteasome (Huang et al., 2015; Suzuki et al., 2016). Recent studies have demonstrated that NGLY1 plays a role beyond ERAD. Specifically, it has been shown that deglycosylation by NGLY1 is required for the translocation and activity of the transcription factor Nrf1 (NFE2L1) (Lehrbach and Ruvkun, 2016; Tomlin et al., 2017). Nrf1 is a key regulator of cellular homeostasis as it controls a broad range of transcriptional programs regulating cellular responses to stress, proteostasis, or metabolic processes (Lee et al., 2011; Lu, 2009; Xu et al., 2005; Zheng et al., 2015).

It is possible that NGLY1 is required for the activity of additional factors besides Nrf1. Recent protein-protein interaction studies report interactions between NGLY1 and various other glycoproteins, some of which may be regulated by NGLY1 (Huttlin et al., 2017, 2020). In addition, the transcriptional output of NGLY1-dependent factors might vary in a cell type dependent manner. As understanding these complex molecular mechanisms might reveal new targets for potential therapeutic interventions, considerable research efforts have been devoted to analyze how NGLY1 deficiency affects cells at the molecular level. Findings from these studies have implicated NGLY1 activity in the regulation of various cellular processes including BMP signaling, mitophagy and the expression of aquaporins (Galeone et al., 2017; Han et al., 2020; Tambe et al., 2019; Yang et al., 2018).

To date, research on NGLY1 deficiency has been conducted predominantly in a variety of model systems including worms, flies, mice, rats and genetically engineered human cell lines (Asahina et al., 2020, 2021; Fujihira et al., 2017, 2020; Galeone et al., 2017; Habibi-Babadi et al., 2010; Mueller et al., 2020; Owings et al., 2018; Rodriguez et al., 2018; Tambe et al., 2019). While these models have proved to be invaluable tools to advance our understanding of NGLY1 biology, it can be challenging to assess how well findings are recapitulated in human patients. To date, however, a systematic analysis of gene and protein expression changes in patient cells has been lacking.

In this study we present the first systematic analysis of gene and protein expression changes in cell lines derived from patients described with NGLY1 deficiency. We used RNA sequencing to measure gene expression in 66 cell lines derived from 14 NGLY1 deficient patients and 17 parental controls. These cell lines represent four different cell types including lymphoblastoid cells (LCLs), fibroblast and fibroblast-derived induced pluripotent stem cells (iPSCs) as well as neuronal progenitor cells. In addition, we performed liquid chromatography coupled with mass spectrometry to obtain protein expression measurements for LCLs and fibroblasts. We show that many previous findings in model organisms are faithfully recapitulated in patient-derived cells. In addition, we identified both cell type-specific and independent dysregulation of processes that had previously not been linked to NGLY1 deficiency. For example, we observed that genes involved in ribosomal mRNA processing were broadly upregulated across cell types. Finally, we provide evidence that gene expression changes in NGLY1 deficient cell lines can be rescued by correcting the mutated alleles using CRISPR/Cas9 (Hsu et al., 2014; Jinek et al., 2012). All results generated in this study can be visualized and explored in an interactive manner at https://apps.embl.de/ngly1browser.

## RESULTS

### Gene expression profiling reveals cell type-agnostic molecular signatures of NGLY1 deficiency

To study how NGLY1 deficiency affects gene expression in patients, we analyzed patient derived cell lines representing four different cell types and measured gene expression by bulk RNA-sequencing (Figure 1a; Supplementary Figure 1a). We first aimed to identify cell type agnostic gene expression changes. We performed a differential gene expression analysis accounting for differences in cell type and sex (Ignatiadis et al., 2016; Love et al., 2014). This analysis revealed 67 significantly downregulated and 39 significantly upregulated genes in NGLY1 deficient cells compared to the control cell lines (FDR < 5%; Supplementary Table 1). Among the top downregulated genes we found NGLY1, VCP, as well as many proteasomal genes. In addition we found previously uncharacterized genes such as FAM219A. Among the most significantly upregulated were genes involved in immune related processes such as NLRP2 or TLR9 (Figure 1b) (Fontalba and Gutierrez, 2007; Hoque et al., 2011).

**Figure 1.**
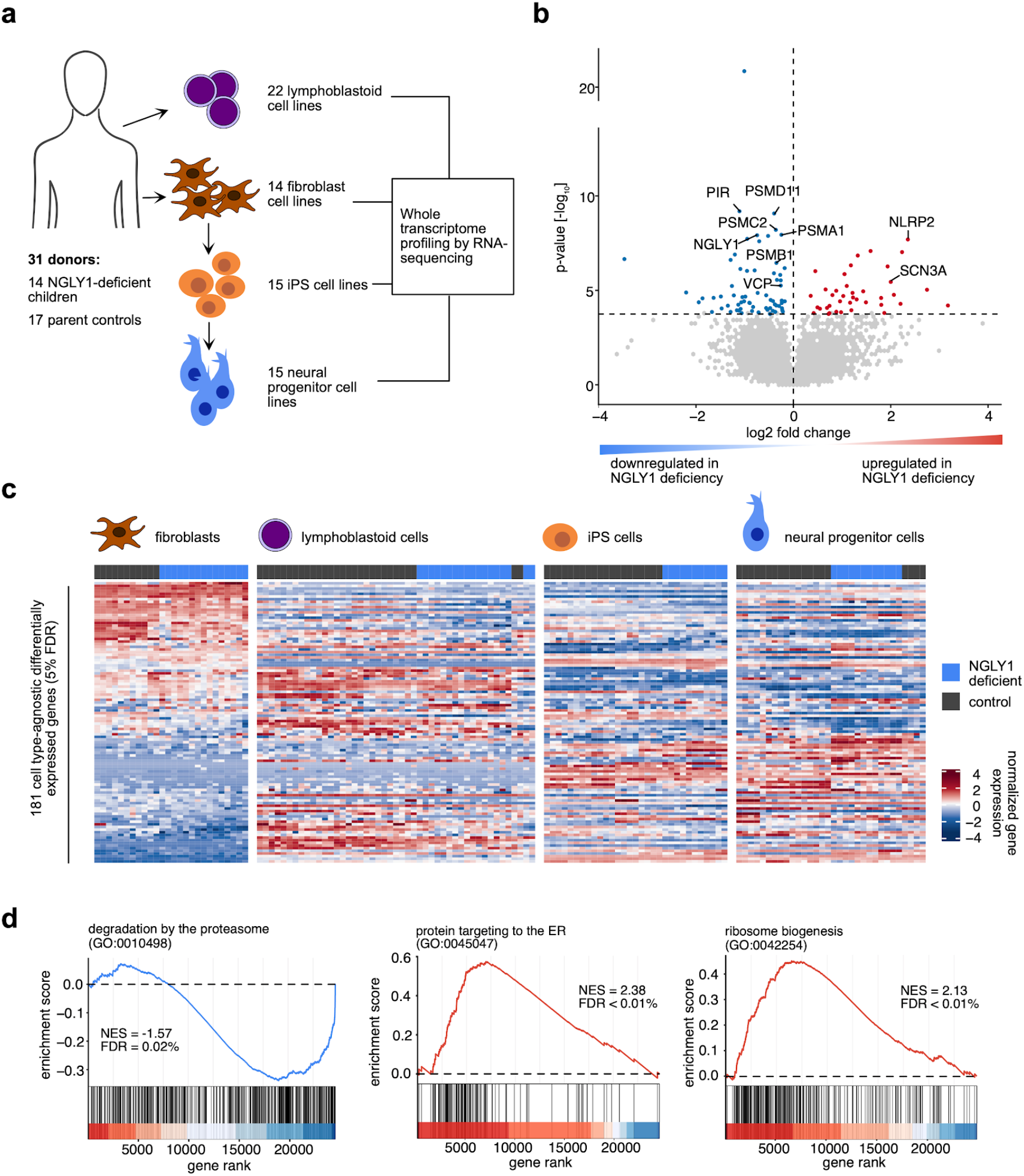
Gene expression profiling reveals cell type-agnostic molecular signatures of NGLY1 deficiency. **(a)** Schematic overview of the panel of patient-derived cell lines used for transcriptome profiling by RNA sequencing. **(b)** Volcano plot of differentially expressed genes in cell lines derived from NGLY1-deficient child patients compared to cell lines derived from parent donors. Blue dots represent genes that are significantly downregulated in NGLY1 deficiency. Red dots correspond to significantly upregulated genes. Selected differentially expressed genes are labeled. The horizontal dashed line marks a false discovery rate (FDR) of 5%. Fold changes and p-values were determined using the DESeq2 R/Bioconductor package. **(c)** Cell lines were grouped by their cell type and clustered based on their standardized expression of 181 genes that were found to be differentially expressed in NGLY1 deficiency. The annotation bar above the heatmaps indicates the NGLY1 deficient cell lines. **(d)** Ranked gene set enrichment analysis results for three selected biological processes in NGLY1 deficient compared to parent control cell lines. Blue and red curves indicate that the corresponding process is down- or upregulated in NGLY1 deficiency, respectively. Enrichment scores and false discovery rates were determined using the clusterProfiler/fgsea R/Bioconductor software. Biological processes are based on Gene Ontology. NES = normalized enrichment score. The complete list of enriched processes can be found in Supplementary Table 2.

Consistent with previous observations in Drosophila, we did not observe an upregulation of genes indicative of increased ER stress (Owings et al., 2018; Samali et al., 2010) (Supplementary Figure 2a). We also did not detect an upregulation of Nrf2 as a potential mechanism to compensate for missing Nrf1 activity (Supplementary Figure 2b). We further failed to detect changes in aquaporin expression (Supplementary Figure 2c), which could indicate that the reported upregulation of aquaporins (Tambe et al., 2019) in NGLY1 deficiency is cell type-specific or can be compensated in patient-derived cells.

To validate the consistency of these differentially regulated genes across cell types we performed a clustering analysis. We clustered all samples corresponding to each individual cell type by their expression of the significantly dysregulated genes (Gu et al., 2016). We found that the samples were consistently grouped by their NGLY1 status in each cell type with few exceptions (Figure 1c). To identify dysregulated pathways in NGLY1 deficient cells we performed a ranked gene set enrichment analysis (Korotkevich et al., 2021; Subramanian et al., 2005; Yu et al., 2012). This revealed 96 significantly upregulated and 88 significantly downregulated processes (Gene Ontology Biological Process terms, FDR < 1%) (The Gene Ontology Consortium, 2019). The enrichment of downregulated processes was primarily driven by the downregulation of proteasomal genes. The upregulated processes could be divided into two functionally related groups represented by an upregulation of tRNA synthesis pathways as well as an upregulation of ribosome biogenesis (Figure 1d, Supplementary Figure 3a). While there was significant upregulation of the immune related genes, such as NLRP2, we did not observe an upregulation of pro-inflammatory gene sets. In addition, the expression of these genes was very low (< 1 TPM) and was not detectable via proteomic analysis.

### Cell type-specific dysregulation of biological processes in NGLY1 deficient cells

We observed that NGLY1 gene expression levels varied between cell types. NGLY1 expression was highest in lymphoblastoid cell lines, while fibroblasts and iPSCs expressed the lowest amount of NGLY1 (Figure 2a). These expression differences might influence how different cell types react to loss of NGLY1. In addition, we hypothesized that NGLY1 might deglycosylate alternative targets in a cell type-specific manner and that Nrf1 target genes might vary between cell types. Therefore, we tested for differentially expressed genes in each cell type separately. We found between ~80 and ~330 genes dysregulated between cell types (5% FDR; Figure 2b). More genes were downregulated than upregulated in all four cell types. Overlap between cell types was surprisingly low (Supplementary Figure 4a). However, this is likely explained at least in part by limited statistical power due to lower sample size and high variability between donors.

**Figure 2.**
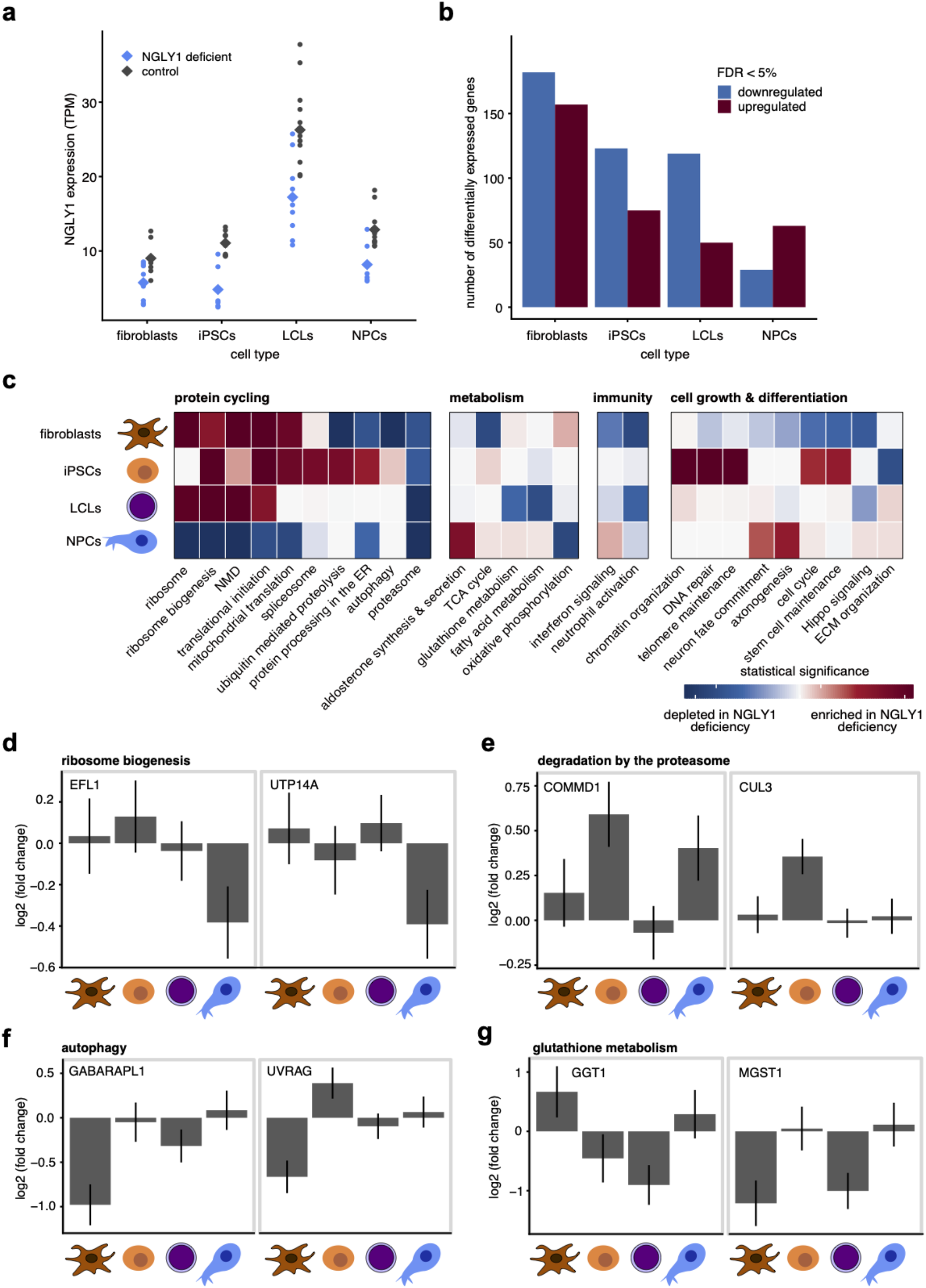
Cell type-specific dysregulation of biological processes in NGLY1 deficient cells. **(a)** Normalized expression of the NGLY1 gene across four different cell types. Each dot represents a patient-derived cell line. NGLY1 deficient cell lines are highlighted in blue. Diamond shaped symbols mark the group means. TPM = transcripts per million; iPSCs = induced pluripotent stem cells; LCLs = lymphoblastoid cell lines; NPCs = neural progenitor cells. **(b)** Number of differentially expressed genes in NGLY1 deficient compared to parent control cell lines. Downregulated and upregulated genes are highlighted in blue and red, respectively. The false discovery rate (FDR) was controlled at 5%. Differential gene expression was determined using DESeq2. **(c)** Ranked gene set enrichment results in different cell types for a list of selected processes. Processes were stratified manually into four biological groups. Each process corresponds to a Gene Ontology Biological Process term, Reactome pathway or KEGG pathway. Colors are based on signed p-values where blue indicates a significant depletion and red indicates a significant enrichment of the process in NGLY1 deficiency. Statistical significance was determined using the clusterProfiler/fgsea R/Bioconductor packages. Detailed statistics can be found in Supplementary Tables 2 and 3. **(d-g)** Examples of cell type specific dysregulated processes in NGLY1 deficiency. For each process the two most strongly dysregulated genes are shown. Fold changes quantify gene expression differences in NGLY1 deficient compared to control cell lines for each cell type. Error bars represent one standard error of the mean and were determined using DESeq2.

For each cell type, we performed ranked gene set enrichment analysis of Gene Ontology Biological Processes, KEGG pathways and Reactome pathways (Jassal et al., 2020; Kanehisa et al., 2021; Korotkevich et al., 2021; Subramanian et al., 2005; The Gene Ontology Consortium, 2019; Yu et al., 2012). Using enrichment maps we identified groups of related processes that were significantly enriched or depleted in each cell type (Merico et al., 2010) (Supplementary Figure 4b). We found overlap but also many cell type-specific effects. For instance, in iPSCs and iPS-derived NPCs we found specific dysregulation of various developmental processes that would not be expected in fibroblasts or LCLs. Figure 2c shows an overview of the effects of NGLY1 deficiency on a number of representative processes. Ribosome and ribosome biogenesis were upregulated in all cell types except for NPCs where we observed the opposite effect (Figure 2d). This downregulation of ribosome biogenesis in NPCs was driven by a set of genes unrelated to those that were upregulated in the other cell types (Figure 2d; Supplementary Table 3). While proteasomal genes were downregulated consistently in all four cell types, we found an upregulation of various ubiquitin ligases in iPSCs (see enrichment of “degradation by the proteasome”, Figure 2e). In addition, we found a fibroblast specific downregulation of autophagy that was driven by a set of genes including GABARAPL1 and UVRAG (Figure 2f). Finally, we found a cell type specific downregulation of genes involved in glutathione metabolism.

In many cases enrichment results were driven by small fold changes. However, these small fold changes were consistent for many genes involved in these processes. This was not only the case for processes previously not linked to NGLY1 deficiency, but also for processes that are well known to be regulated by NGLY1 and its substrate Nrf1, such as the proteasome (Tomlin et al., 2017) (Supplementary Figure 4c).

### Gene expression changes in NGLY1 deficient cells can be reverted by correcting NGLY1 mutations using CRISPR-Cas9

The majority of patients described with NGLY1 deficiency are less than 20 years old. The parents of these patients typically volunteer as controls. The resulting age difference between disease cases and controls can make it difficult to separate disease-from age associated effects. We asked whether the gene expression differences we observed were in fact due to NGLY1 deficiency. In addition, we aimed to explore whether these changes could be reverted by correcting the mutated NGLY1 alleles. To answer these questions, we obtained iPS and neural progenitor cell lines derived from one patient donor (CP1) where NGLY1 mutations in exons 8 and 11 had been corrected using CRISPR-Cas9 (Hsu et al., 2014; Jinek et al., 2012). To account for potential clonality effects we obtained two independent clones per correction (12 cell lines in total, 6 per cell type). We then performed bulk RNA sequencing and processed the data as described. For both cell types we performed two independent differential gene expression analyses and compared gene expression in the NGLY1 deficient cells to either the parent control cell lines or the CRISPR-corrected cell lines.

In iPSCs the results agreed well between the tests that were performed using different types of controls (R = 0.47, Figure 3a). As expected, we observed that proteasomal genes were downregulated in NGLY1 deficient cell lines compared to both parent and CRISPR-corrected controls. The upregulation of ubiquitin ligases such as CUL3 or COMMD1 was also confirmed by a rescue effect in the corrected cells (i.e. their expression decreased upon CRISPR-correction of patient NGLY1 mutations). Interestingly, we also found that the downregulation of the uncharacterized gene FAM219A was rescued when NGLY1 mutations were reverted (Figure 3b). Looking at the neural progenitor cell lines we found that the overlap between the differential gene expression results was less pronounced but still significant (R = 0.22, p < 2.2e-16; Figure 3c). Similar to the iPSCs we observed that the downregulation of proteasomal genes and FAM219A was reverted in the NGLY1-corrected cell lines. We observed that transcription factors important for neuronal development, such as OLIG2 (Zhou and Anderson, 2002; Zhou et al., 2001), were upregulated in NGLY1 deficient neural progenitor cells (Figure 3d). Despite the fact that these genes are expressed only at very low levels (< 1 TPM) in our cell lines, their upregulation was reverted in the CRISPR corrected lines adding credibility to this observation. When comparing NGLY1 deficient cell lines and cell lines derived from parent donors, we found that important stem cell factors, such as LIN28A, POU5F1 (Oct4) or NANOG (Takahashi and Yamanaka, 2006; Takahashi et al., 2007; Viswanathan and Daley, 2010), were downregulated in NGLY1 deficient cells. This downregulation, however, was not reverted in the CRISPR corrected cell lines where these factors were expressed at similar levels as in the NGLY1 deficient lines. This suggests that the differential expression of stem cell factors is explained by a difference in donor age rather than the lack of NGLY1. In summary, we observe that key gene expression differences between patient and parent control cell lines can be explained by a lack of NGLY1 although age-associated differences can confound these results in a cell type-specific manner.

**Figure 3.**
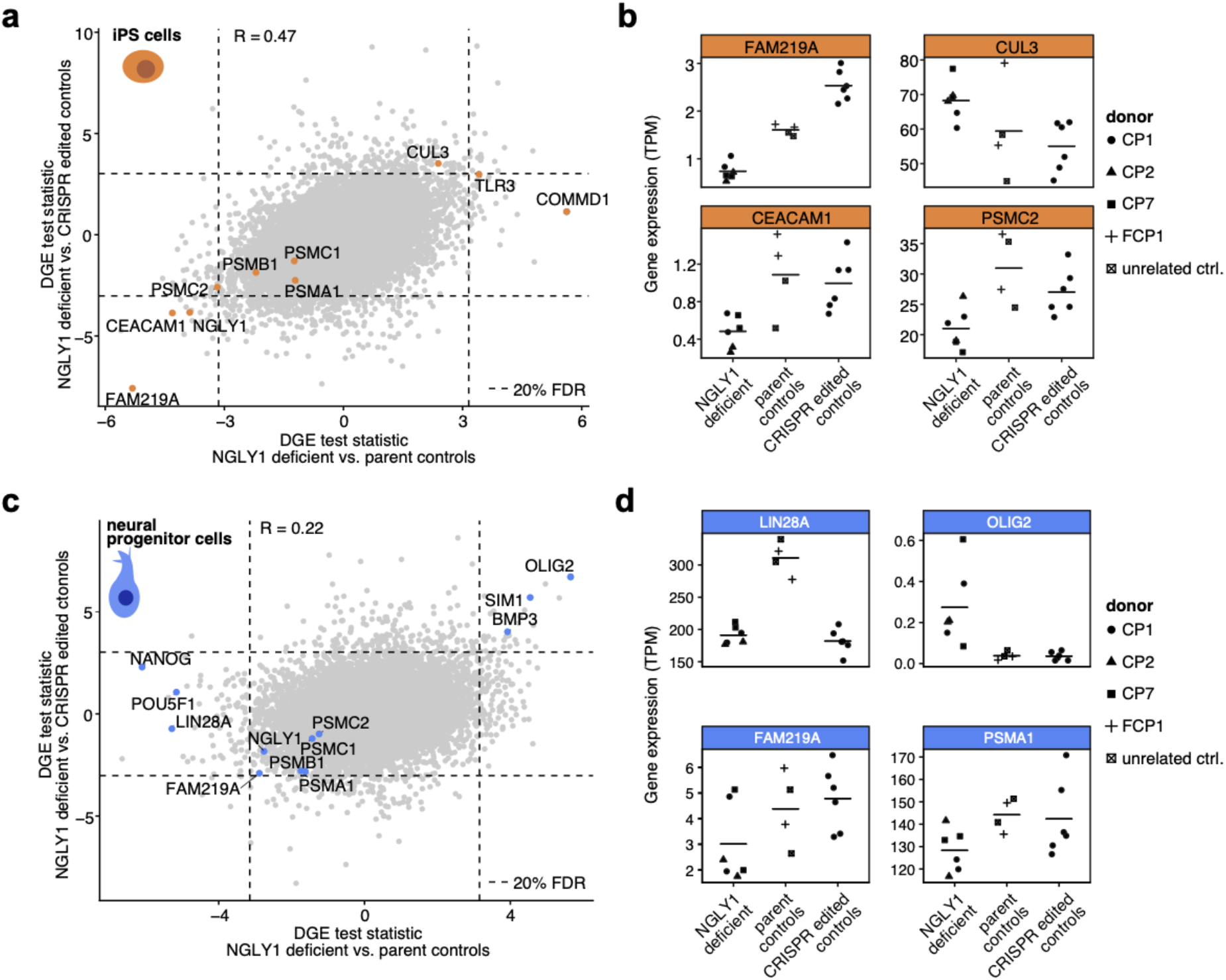
Gene expression changes in NGLY1 deficient cells can be reverted by correcting NGLY1 mutations using CRISPR-Cas9. **(a)** Differential gene expression in NGLY1 deficient induced pluripotent stem cells was determined by comparing patient cell lines to parent controls (x-axis) or to patient cell lines where the NGLY1 allele had been corrected using CRISPR/Cas9 (y-axis). Horizontal and vertical dashed lines indicate a false discovery rate of 20%. Selected genes are labeled and highlighted in orange. Differential gene expression was determined using DESeq2. R = Pearson’s correlation coefficient. **(b)** Normalized gene expression values for four selected genes in NGLY1 deficient induced pluripotent stem cell lines, parent controls and CRISPR/Cas9 corrected controls. Different symbols identify cell lines derived from different donors. Horizontal lines mark the group averages. **(c)** Differentially expressed genes in NGLY1 deficient neural progenitor cell lines compared to parent cell lines and CRISPR/Cas9 corrected controls. Selected genes are highlighted in blue and labeled. **(d)** Normalized gene expression values for four selected genes in NGLY1 deficient neural progenitor cell lines, parent controls and CRISPR/Cas9 corrected controls.

### High-throughput proteomic analysis reveals downregulation of ER-associated degradation and glutathione biosynthesis

NGLY1 functions as part of the endoplasmic reticulum associated degradation pathway and thus acts as a regulator of protein turnover (Christianson et al., 2011). Accordingly, we expected that NGLY1 deficiency might lead to changes in protein levels, even when mRNA levels are not affected. To examine the effects of NGLY1 deficiency on protein expression, we applied liquid chromatography coupled with mass spectrometry to systematically measure protein levels in our patient-derived fibroblast and LCL cell lines (Figure 4a).

**Figure 4.**
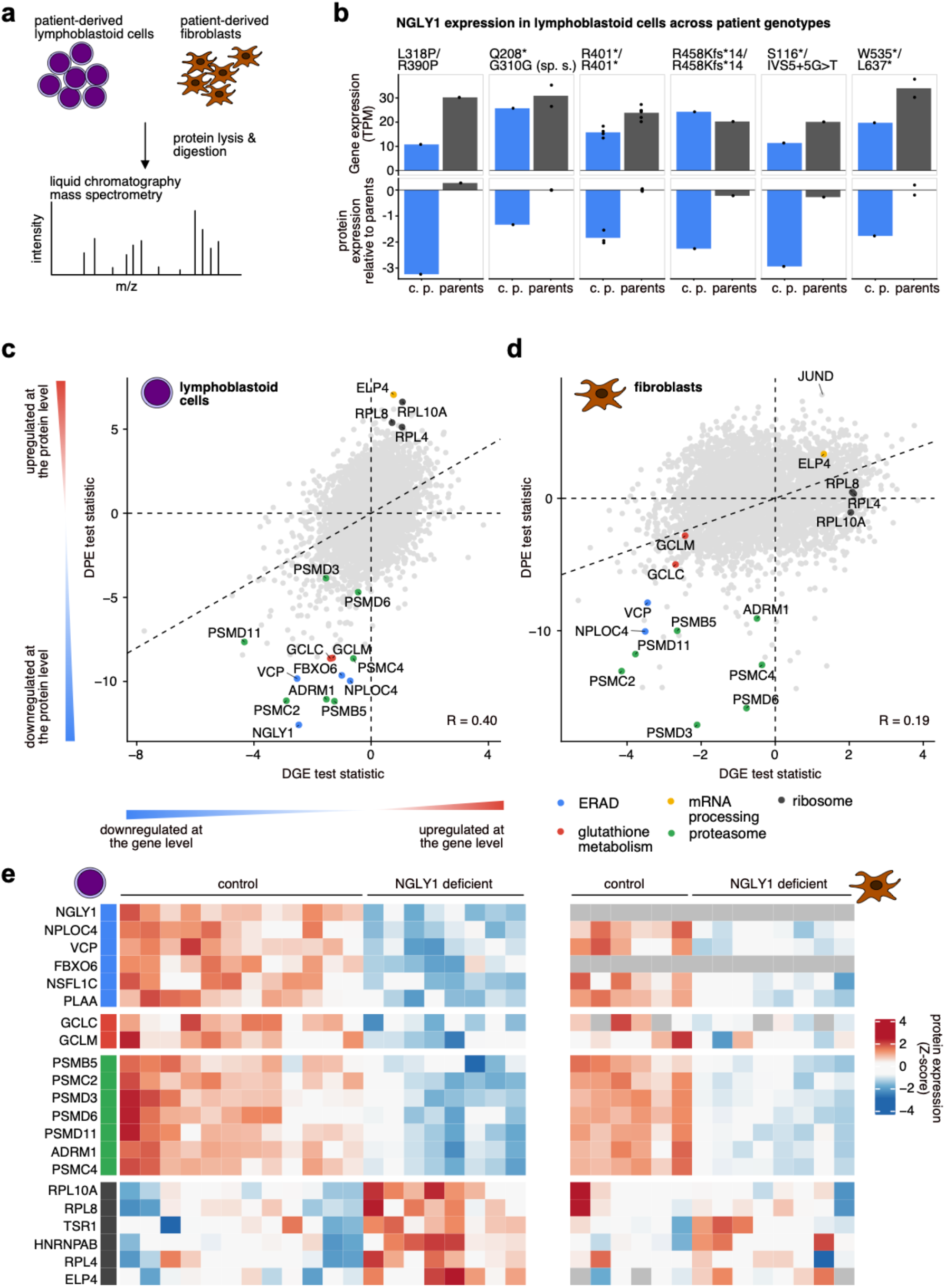
High-throughput proteomic analysis reveals downregulation of ER-associated degradation and glutathione biosynthesis. **(a)** The same lymphoblastoid and fibroblast cell lines that were analyzed using RNA sequencing were subjected to liquid chromatography mass spectrometry to measure protein expression. **(b)** NGLY1 gene (top) and protein (bottom) expression across different patient genotypes. For each genotype NGLY1 expression is compared between the corresponding patients and their parents. Each dot corresponds to one cell line and the bars represent the average expression per donor group. Normalized protein expression was determined using the DEP R/Bioconductor package. TPM = transcripts per million. **(c-d)** Differential gene expression (x-axis) and differential protein expression (y-axis) statistics in NGLY1 deficient lymphoblastoid (c) or fibroblast (d) cell lines compared to parent controls. Only genes that were detected by both RNA sequencing and mass spectrometry are shown. Selected genes are labeled and colored according to their associated biological processes. Differential gene expression statistics were determined using DESeq2. Differential protein expression was inferred using a linear mixed effects model. R = Pearson’s correlation coefficient; ERAD = endoplasmic-reticulum-associated degradation. **(e)** Heatmap view of selected differentially expressed proteins in NGLY1 deficient lymphoblastoid cells (left) and fibroblasts (right). The selected genes (rows) are part of four different biological processes and are grouped accordingly. Cell lines (columns) are grouped into NGLY1 deficient and control cell lines. Grey tiles in the heatmap indicate that a protein could not be detected in the corresponding cell line.

We first analyzed NGLY1 protein expression across samples. While NGLY1 protein could be detected in all lymphoblastoid cell lines, we were unable to detect NGLY1 expression in any of the fibroblastoid cell lines independent of the donor. Since NGLY1 gene expression is much lower in fibroblasts compared to LCLs (Figure 2a) it is possible that protein expression of NGLY1 in fibroblasts is below the detection limit of our assay. We compared NGLY1 protein expression in patient-derived lymphoblastoid cell lines to the parent-derived cell lines. We observed a 2.5-9.5 fold downregulation of NGLY1 protein expression in patients compared to parent-derived cell lines depending on the patient genotype (Figure 4b). Comparing NGLY1 protein expression to the expression of all other detected proteins in LCLs we observed that NGLY1 was among the 0.2%-11% most lowly expressed proteins in patient-derived cell lines. In contrast, NGLY1 was among the 24%-45% most lowly expressed proteins in parent controls and among the 46%-71% most lowly expressed proteins in a cell line derived from an unrelated control subject (Supplementary Figure 5a-b).

We tested for differential protein expression in both LCLs and fibroblasts. In total we found that 1527 and 679 proteins were significantly down- or upregulated in fibroblasts and LCLs, respectively (FDR < 5%; Supplementary Table 4). Similar to the gene expression analysis, we found proteasomal subunits, proteins involved in ER-associated degradation and proteins associated with ribosomal mRNA processing among the most significantly dysregulated proteins in both cell lines (Figure 4c-d). Overall, differential gene and protein expression analyses agreed well with each other (R = 0.4 for LCLs and R = 0.19 for fibroblasts, Figure 4a-b) although in many cases, however, we found that differential expression was more pronounced and thus easier to detect at the protein level compared to the gene level (Figure 4a-b). This enabled us to detect protein expression changes that we had not been able to observe at the transcript level. For example, we found several proteins involved in ER associated degradation, such as FBXO6 or NPLOC4 (Bays et al., 2001; Yoshida et al., 2003), to be downregulated in the NGLY1 deficient cell lines. In addition we observed an LCL-specific downregulation of GCLC and GCLM, two subunits of the glutamate-cystein ligase which catalyzes a rate limiting step in the production of glutathione and which has previously been shown to be regulated by Nrf1 (Myhrstad et al., 2001; Yang et al., 2005). While we did not find this specific enzyme to be downregulated at the transcript level, this observation agrees with our cell-type specific gene set enrichment analysis, which had shown a general LCL-specific downregulation of genes involved in glutathione metabolism (Figure 2c). The dysregulation of proteins involved in ER associated proteasomal degradation, glutathione synthesis and ribosomal mRNA processing were consistent across individual donors, suggesting that these changes are not specific to certain patient subgroups (Figure 4e).

### A web application enables community access to gene expression and proteomic signatures of NGLY1 deficiency

In recent years, many key contributions towards a better understanding of NGLY1 deficiency have been made thanks to research in model organisms. Generally, findings in model systems have to be assessed for their relevance to human disease. To facilitate this task, we developed a web application that allows users to quickly look up whether a gene of interest is differentially expressed in cell lines derived from NGLY1 deficient patients. This online tool is available at https://apps.embl.de/ngly1browser and is built upon a database that contains all data and analysis results generated in this study. The application consists of four panels. The first two panels visualize mRNA and protein expression of individual genes of interest in different cell types (Figure 5a). After a user selects gene and cell type, a bar plot shows normalized expression values across all relevant samples in the database. In addition, a scatter plot shows summarized statistics for patient and control samples. A third panel allows users to enter a list of gene identifiers and examine if these genes are dysregulated in NGLY1 deficiency in comparison to other genes. To achieve this volcano plots that show fold changes and p-values are displayed for various cell types. Finally, a fourth panel in the web application is dedicated to gene set enrichment analysis. Here, the user can specify cell type, experiment type and one of currently three gene set databases (Gene Ontology, Reactome and KEGG) (Jassal et al., 2020; Kanehisa et al., 2021; The Gene Ontology Consortium, 2019). Processes that are significantly enriched or depleted in patient derived cell lines are displayed as a graph, where each node represents a gene set and connections between the nodes indicate that the gene sets consist of similar genes (Figure 5b). Users can click on nodes in order to see the genes that drive the enrichment of its corresponding pathway. For each gene, detailed statistics regarding their differential expression in NGLY1 deficiency will be shown in tabular format. We envision that this web application and database will be of broad use to the NGLY1 research community. In the future, we aim to further extend the database by including additional experiments in new cell types.

**Figure 5.**
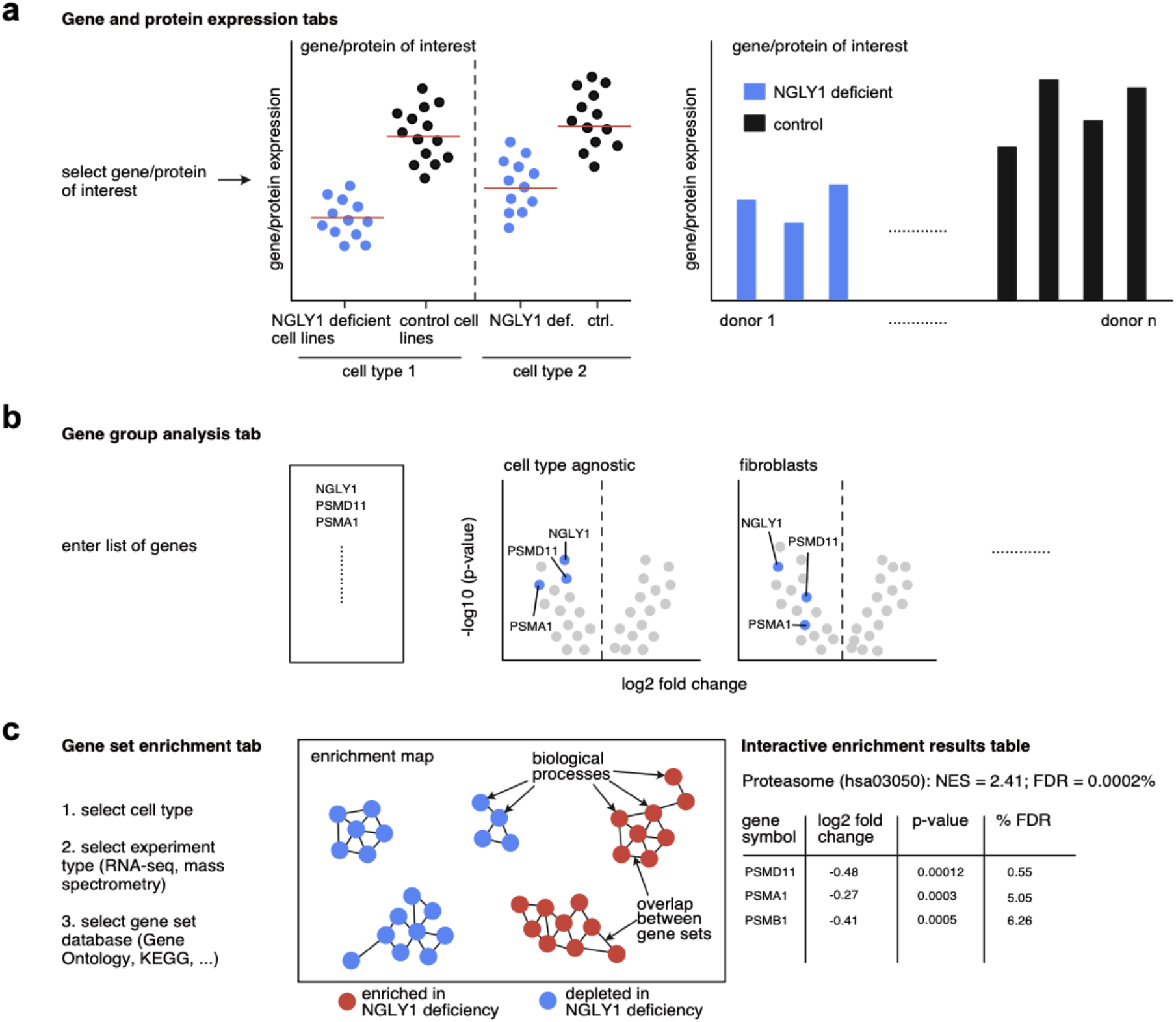
A web application enables community access to gene expression and proteomic signatures of NGLY1 deficiency. **(a)** Gene and protein expression tabs in the NGLY1 browser web application allow users to select a gene of interest and visualize its expression across all samples in the database. Expression values are shown for each cell type separately. In addition, a scatter plot summarizes the expression values by NGLY1 deficiency status to allow for easy assessment of disease specific expression differences. **(b)** A gene group analysis tab allows users to enter a list of genes and examine their differential expression in NGLY1 deficiency. For each gene, log2 fold changes and p-values are visualized in the context of all other genes as volcano plots. Results are available for both cell type-specific and agnostic analyses. **(c)** A gene set enrichment analysis tab shows significantly enriched or depleted biological pathways from different gene set databases as a graph in which each node represents a pathway. Nodes are colored depending on whether the corresponding pathway is up- or downregulated in NGLY1 deficiency. Nodes are connected by edges if the enrichment of their associated biological processes is driven by similar genes. Clicking on the nodes displays additional information in tabular format.

## DISCUSSION

Using RNA sequencing and mass spectrometry we generated the first map of cell type-specific gene and protein expression changes in NGLY1 deficient patients. This map provides a resource to guide future experiments in model systems in directions most relevant to human disease. We have made all analysis results presented in this study available at https://apps.embl.de/ngly1browser. In the future we aim to integrate new datasets into this resource as they become available.

Many of the gene expression changes we observed in NGLY1 deficient cells were subtle. This was not only the case for genes not previously linked to NGLY1 deficiency but also for genes that have long been known to be regulated by NGLY1. To convincingly identify small changes in gene expression a substantial number of samples is required - especially when the variability between individual donors is large. This poses a significant challenge when dealing with a rare disease. While fold changes were often small, we observed that dysregulation was often very consistent across multiple genes that are part of the same biological process. We took advantage of this by relying on gene set enrichment analysis so as to obtain sufficient statistical power to identify dysregulated gene groups. It is possible, however, that small but biologically significant expression changes of individual genes or insufficiently annotated pathways were missed and identifying these will require additional samples.

We observed that many of the gene expression changes in NGLY1 deficiency were cell type-specific. It is likely that due to small sample sizes, this non-overlap is overestimated. For example, some proteasomal genes could not be detected as significantly downregulated in some cell types due to small fold changes and high donor-to-donor variability. Nevertheless, it is likely that many of the cell type-specific effects we observe are in fact real. For instance, at the proteomics level the downregulation of GCLC and GCLM, two genes known to be regulated by Nrf1 (Myhrstad et al., 2001; Yang et al., 2005), was very prominent in LCLs but completely absent in fibroblasts. Related to previous findings in *C. elegans* (Habibi-Babadi et al., 2010), we further observed a dysregulation of important neural transcription factors in NPCs that could be rescued by correcting NGLY1 mutations via CRISPR/Cas9. These observations suggest that NGLY1 might have additional effects on gene and protein expression in other cell types that have not yet been examined. Since some of these might be druggable it could be worthwhile to perform experiments in additional cell types.

We used mass spectrometry to measure protein expression in NGLY1 deficient cell lines. This revealed that in patient LCLs NGLY1 protein was, although strongly downregulated, still detectable in all cell lines. This could be a positive sign. A potential approach to combat NGLY1 deficiency is to re-establish NGLY1 expression by gene therapy, gene correction, or enzyme replacement therapy. For such therapies, potential immune reactions to the replacement protein are often a concern. The fact that some remaining NGLY1 protein is still present naturally in the patient cell lines might indicate that such an immune reaction is less likely to occur.

NGLY1 encodes a deglycosylation enzyme so the absence of functional NGLY1 protein is likely to impact the post-translational state of its targets. Using mass spectrometry we measured the expression of proteins across patient-derived cell lines. However, these experiments did not allow us to assess the posttranslational modifications (PTMs) of these proteins. In the future, measurements of PTMs might yield additional insights into how NGLY1 deficiency affects protein function. Further, we focused on measuring gene and protein expression in entire cells. Examining and comparing protein abundance in specific cell fractions or under specific cellular stress conditions might reveal additional details of how NGLY1 deficiency affects protein function across cellular compartments and conditions.

Our results demonstrate that many previous findings on NGLY1 deficiency that were made in model organisms are recapitulated in patient-derived cell lines. In addition, we report gene and protein expression changes in various pathways that had previously not been linked to NGLY1 and whose biological and clinical significance remains to be evaluated. These observations will guide future research towards a more complete understanding of NGLY1 deficiency and how it can be overcome.

## METHODS

### Cell culture

Lymphoblastoid cell lines were maintained in RPMI1640 with 10% FCS. Fibroblast cell lines were maintained in DMEM with 10% FCS. Lymphoblastoid and fibroblast cell lines are available through Coriel by searching using the designation “CONGENITAL DISORDER OF DEGLYCOSYLATION; CDDGN-GLYCANASE 1; NGLY1”.

### Generation of iPSCs and NPCs

Patient’s skin biopsy samples were collected in Stanford Hospital and fibroblast cells were derived in Stanford Cytogenetics Laboratory. The iPSCs were generated with the Sendaivirus delivery system (Fusaki et al., 2009) using the CytoTune-iPS 2.0 Sendai Reprogramming Kit (ThermoFisher Scientific). Briefly, the fibroblast cells were maintained in DMEM containing 10% fetal bovine serum. During the reprogramming, 2×10^5^ cells were transduced with Sendai virus with MOI of 5:5:3 (KOS:c-Myc:Klf4) as instructed in the manual. The cells were maintained in fibroblast media for 6 days then passaged onto Matrigel-coated dishes for further reprogramming in Essential 6 medium plus human bFGF (ThermoFisher Scientific). The iPSC colonies were manually picked onto Matrigel-coated plates to generate stable iPSC lines. The iPSCs were maintained in mTeSR1 medium (Stemcell Technologies) and routinely passaged every 4~6 days with Versene (EDTA) solution (Lonza). Selected iPSC clones were further validated for expression of pluripotency markers with immunostaining (Chen et al., in preparation) and NGLY1 mutations were confirmed with Sanger sequencing. The karyotyping of the iPSCs were checked by G-banding analysis at the WiCell institute.

### Feeder-free differentiation of iPSCs to NPCs

Feeder-free NPC differentiation was achieved by using a modified version of the dual-SMAD signaling inhibition approach previously published in Chambers *et al* (Chambers et al., 2009). iPSC monolayer cultures were incubated in KO-DMEM medium containing 20% knockout serum replacement (KSR, ThermoFisher) with 10uM SB431542, 100nM LDN-193189, Glutamax (1/100), NEAA (1/100) and beta-mercaptoethanol (0.1mM). The cells were fed with fresh media every other day and collected seven days after differentiation.

### RNA extraction and RNA sequencing library preparation

RNA was extracted with Trizol according to the manufacturer’s instructions. This RNA was used to create libraries for RNA-sequencing using the Illumina TruSeq Kit protocol according to the manufacturer’s specifications. RNA sequencing was performed on an Illumina NextSeq machine to generate 75 bp single end reads.

### Analysis of RNA sequencing data

RNA-seq reads were quantified against the human genome version GRCh38 (Stolarczyk et al., 2020) using the software Salmon v.1.2.0 in alignment free mode (Patro et al., 2017). For quality control, a variance stabilizing transformation was applied to the sample read counts as implemented in the DESeq2 R/Bioconductor package (Huber et al., 2015; Love et al., 2014). Clustering as well as principal component analysis were used to identify technical outliers (Gu et al., 2016). In total, 4 out of 140 total samples were excluded from downstream analysis (Supplementary Figure 1). Technical replicates were merged using the ‘collapseReplicates’ function implemented in DESeq2 (Love et al., 2014). Both uncorrected and CRISPR corrected cell lines were used for differential gene expression analysis of NGLY1 deficiency. Cell lines derived from parent or unrelated donors as well as patient-derived cell lines with at least one corrected NGLY1 allele were labeled as ‘NGLY1 functional’. All uncorrected patient-derived cell lines were labeled ‘NGLY1 deficient’. To identify cell-type agnostic differentially expressed genes DESeq2 was used accounting for differences in donor sex and cell type (Love et al., 2014). Shrinkage of log2 fold changes was performed using the ‘apeglm’ method (Zhu et al., 2019). For analysis of cell type-specific differentially expressed genes, samples were grouped according to their cell type and NGLY1 status. A beta prior and a local dispersion fit were applied when running DESeq2. The false discovery rate of differentially expressed genes was controlled at 5% using the Benjamini-Hochberg method.

### Proteomic sample processing and LC-MS/MS analysis

Samples were pre-fractionated with an off-line UltiMate 3000 LPG LC system (Thermo Fisher Scientific), using a basic pH reverse phase separation. Whole cell lysates were fractionated and pooled into 25 fractions. Of these, initially 11 fractions were measured over 120 min on a reverse phase LC gradient, online-injected into a Q Exactive MS instrument (Thermo Fisher Scientific), and data were generated for MS2 applying top10, HCD fragmentation, peptide matching, exclusion of isotopes and dynamic exclusion of precursors. TMT reporter and peptide fragment (amino acid sequence) information was generated in one spectrum and calculated/analyzed/reported by an in-house written software. Database search was done using a Mascot server and the human IPI database. Analysis was carried out on a Q Exactive Plus or Q Exactive HF (both Thermo Fisher Scientific) mass spectrometers coupled to UltiMate 3000 RSLC Nano LC systems (Thermo Fisher Scientific).

Raw mass spectra were analyzed by Mascot and IsobarQuant (Franken et al., 2015). Normalization of raw mass spectra was performed using the DEP R/Bioconductor package (Zhang et al., 2018). To identify differentially expressed proteins, a linear mixed-effects model of the form *y_p,i_* = a_*j*[*p,i*]_ + *X_i_β* + *ε_p i_* was fit to the data using the R package ‘lme4’ (Bates et al., 2014). Here *y_p,i_* is the normalized protein expression of protein *p* in sample *i, α*_*j*[*p,i*]_ represents the baseline protein expression in experimental batch *j* and *X_i_β* represents the NGLY1 deficiency status of sample *i*. *ε_p,i_* is a noise term that models biological and experimental variation. Independent models were fit for each cell type to identify cell typespecific protein changes. Statistical significance of mixed effects model coefficients was estimated using the ‘lmerTest’ R package (Kuznetsova et al., 2017). Test statistics estimated by ‘lme4’ were used to compare protein expression and gene expression changes (where test statistics were estimated by DESeq2 (Love et al., 2014)). The false discovery rate was controlled at 5% using the Benjamini-Hochberg method.

### Gene set enrichment analysis

Ranked gene set enrichment analysis of Gene Ontology biological processes (The Gene Ontology Consortium, 2019), Reactome pathways (Jassal et al., 2020) and KEGG pathways (Kanehisa et al., 2021) was performed using the ‘fgsea’ method as implemented in the clusterProfiler R/Bioconductor package (Huber et al., 2015; Korotkevich et al., 2021; Yu et al., 2012). The test statistics generated by DESeq2 (for RNA-seq data) or the linear mixed effects model (for mass spectrometry data) were used to rank genes. The false discovery rate of enriched pathways was controlled at 1% using the Benjamini-Hochberg method. Significantly enriched processes were grouped based on the number of shared genes by generating an enrichment map for each individual GSEA test (Merico et al., 2010; Yu, 2018; Yu et al., 2012). To generate Figure 2c a representative term was selected manually for each cluster of terms in the enrichment maps (Supplementary Table 3, Supplementary Figure 4b).

## ACKNOWLEDGEMENTS

We are grateful to William A. Gahl, M.D., Ph.D. and Lynne Wolfe, MS, CRNP, BC for sharing patient cell lines. This work was funded by the Grace Science Foundation.

## COMPETING INTERESTS

The authors declare no conflict of interest.

## Notes

https://apps.embl.de/ngly1browser

### Competing Interest Statement

The authors have declared no competing interest.

