## Supplementary Figures for "Patient-derived gene and protein expression signatures of NGLY1 deficiency"

**a**

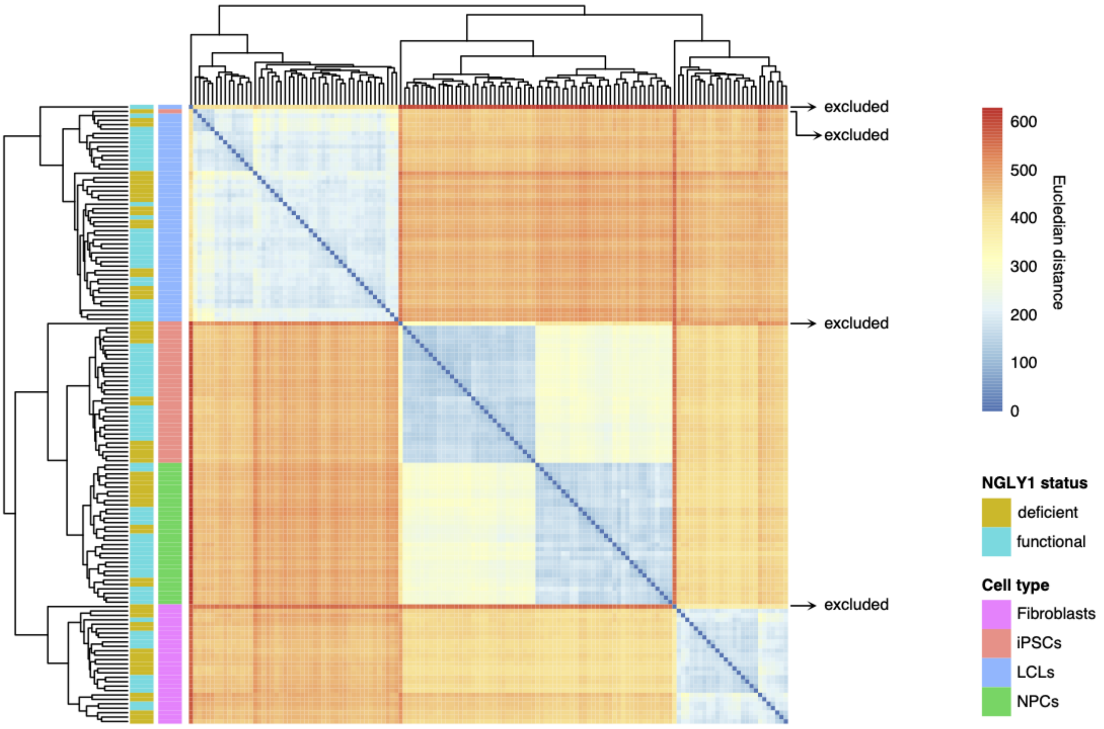


**Supplementary Figure 1.** Hierarchical clustering of bulk RNA-seq datasets analyzed in the study. **(a)** A heatmap based on the Euclidean distance between all sample pairs. The distance was calculated based on normalized gene expression values after applying a variance stabilizing transformation as implemented in the DESeq2 R package. Sample clustering was performed based on the complete linkage method. The annotation bars on the left indicate NGLY1 status and cell type. The NGLY1 status "functional" includes both samples with no mutation of NGLY1 and samples with heterozygous NGLY1 mutations. Arrows indicate outlier samples that were removed from downstream analyses.

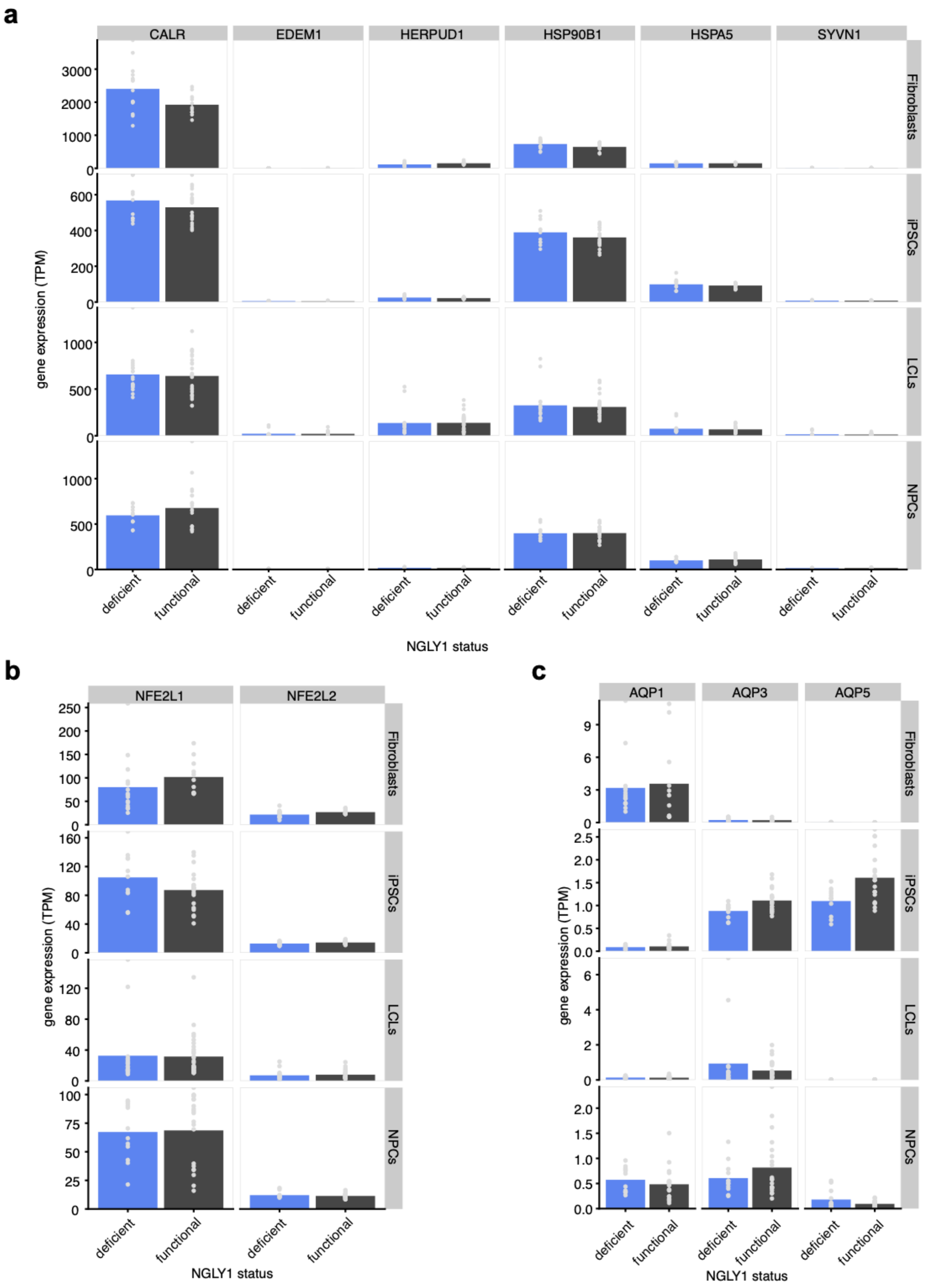


**Supplementary Figure 2.** Markers of ER stress, Nrf1/2 and aquaporins are not differentially expressed in the patient-derived cell lines analyzed in this study. **(a)** Expression of marker genes that are typically upregulated under ER stress (based on Samali *et al.*, 2010). **(b)** Expression of the transcription factors Nrf1 (NFE2L1) and Nrf2 (NFE2L2). **(c)** Expression of selected aquaporins previously shown to be upregulated in NGLY1 deficient mouse embryonic fibroblasts. Bars represent group means. Grey dots indicate gene expression in individual samples. The NGLY1 status “functional” includes both samples with no mutation of NGLY1 and samples with heterozygous NGLY1 mutations (parent controls). TPM = transcripts per million.

**a**

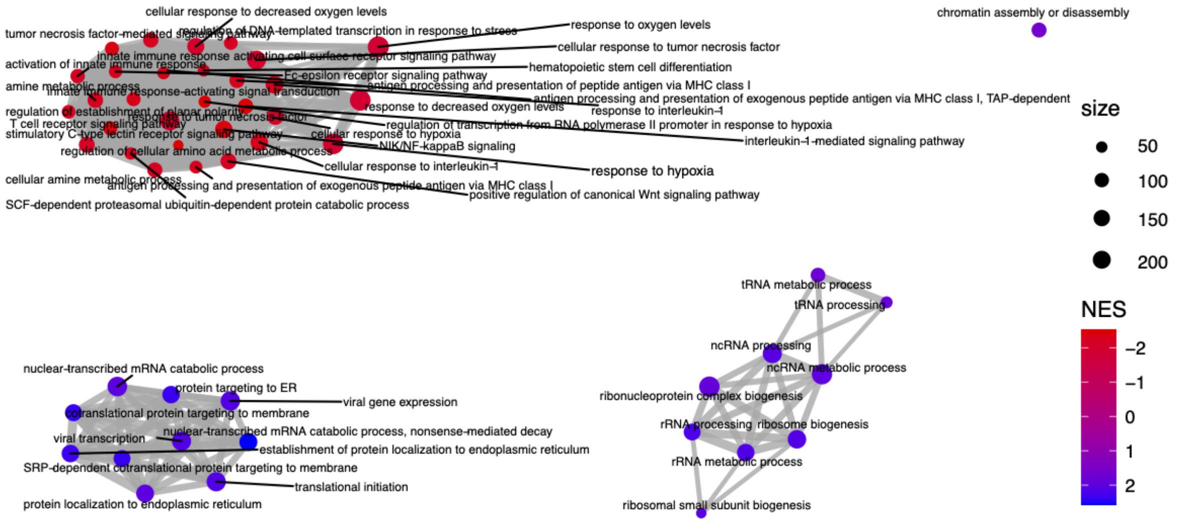


**Supplementary Figure 3.** Enrichment of Gene Ontology (GO) Biological Process terms in NGLY1 deficient cells (cell type-agnostic analysis). **(a)** Enrichment map of the 30 most upregulated (blue) and the 30 most downregulated (red) GO Biological Process terms based on a ranked gene set enrichment analysis (FDR < 0.1% in all cases). Each node in the network represents a GO term, grey links between nodes indicate an overlap between the genes associated with each GO term. Larger nodes indicate that a higher number of genes is associated with that term. Thicker links indicate that a larger number of genes is shared between two terms. NES = normalized enrichment score.

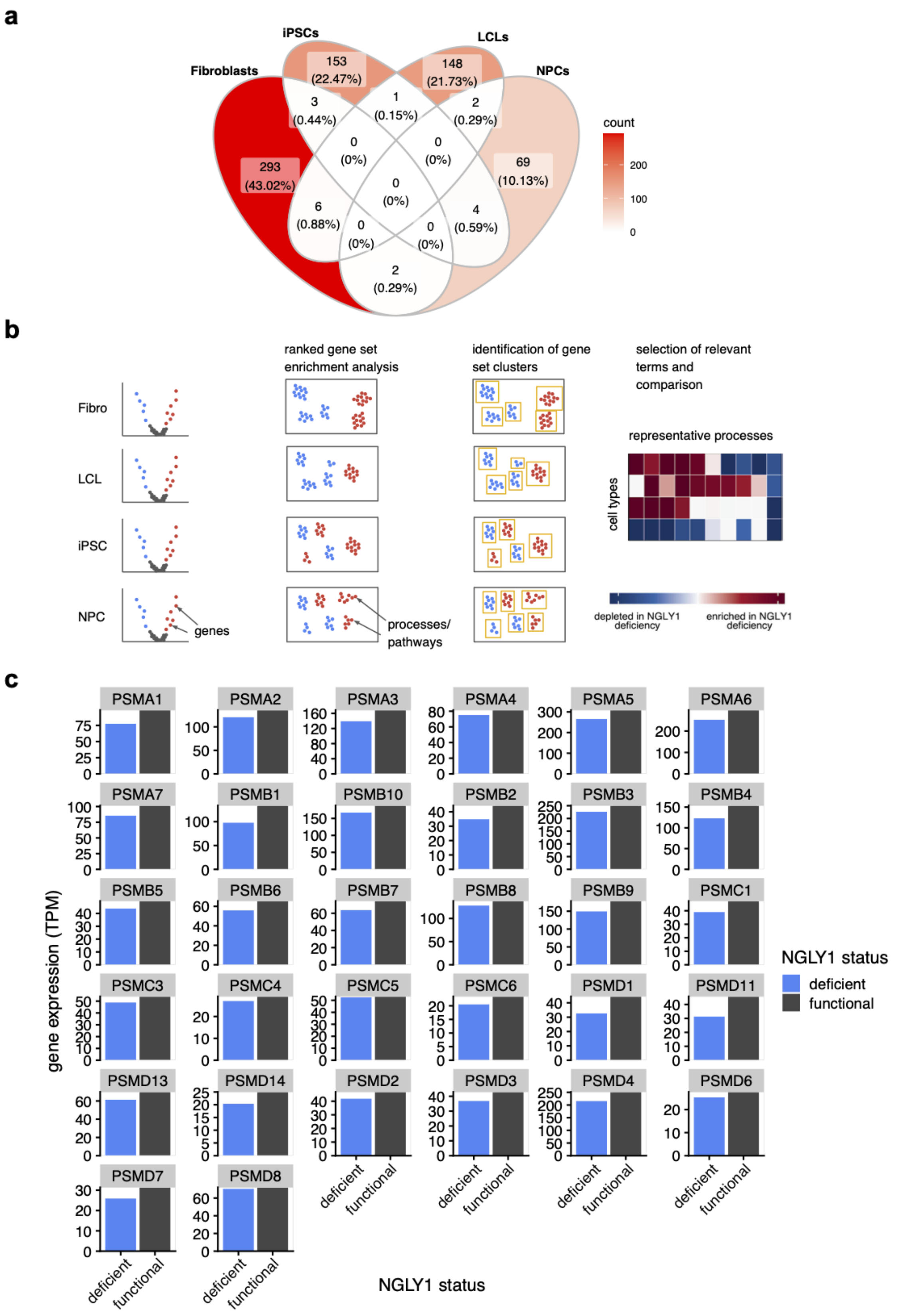


**Supplementary Figure 4. (a)** Venn diagram of differentially expressed gene sets (FDR < 5%) in each NGLY1 deficient cell type. Red color indicates larger set size. **(b)** Schematic depiction of the enrichment analysis process to generate Figure 2c. DESeq2 was used to rank genes based on their differential expression in each NGLY1 deficient cell type (based on the DGE test statistic). A ranked gene set enrichment analysis of GO Biological Processes, Reactome pathways and KEGG pathways was performed for each list of DGEs and an enrichment map was generated. A representative term was selected manually for clusters in the enrichment maps and these terms were combined in a heatmap. **(c)** Expression of proteasomal genes (20s and 26s proteasome subunits) in NGLY1 deficient cells compared to controls in lymphoblastoid cell lines (LCLs). The NGLY1 status “functional” includes both samples with no mutation of NGLY1 and samples with heterozygous NGLY1 mutations (parent controls). TPM = transcripts per million.

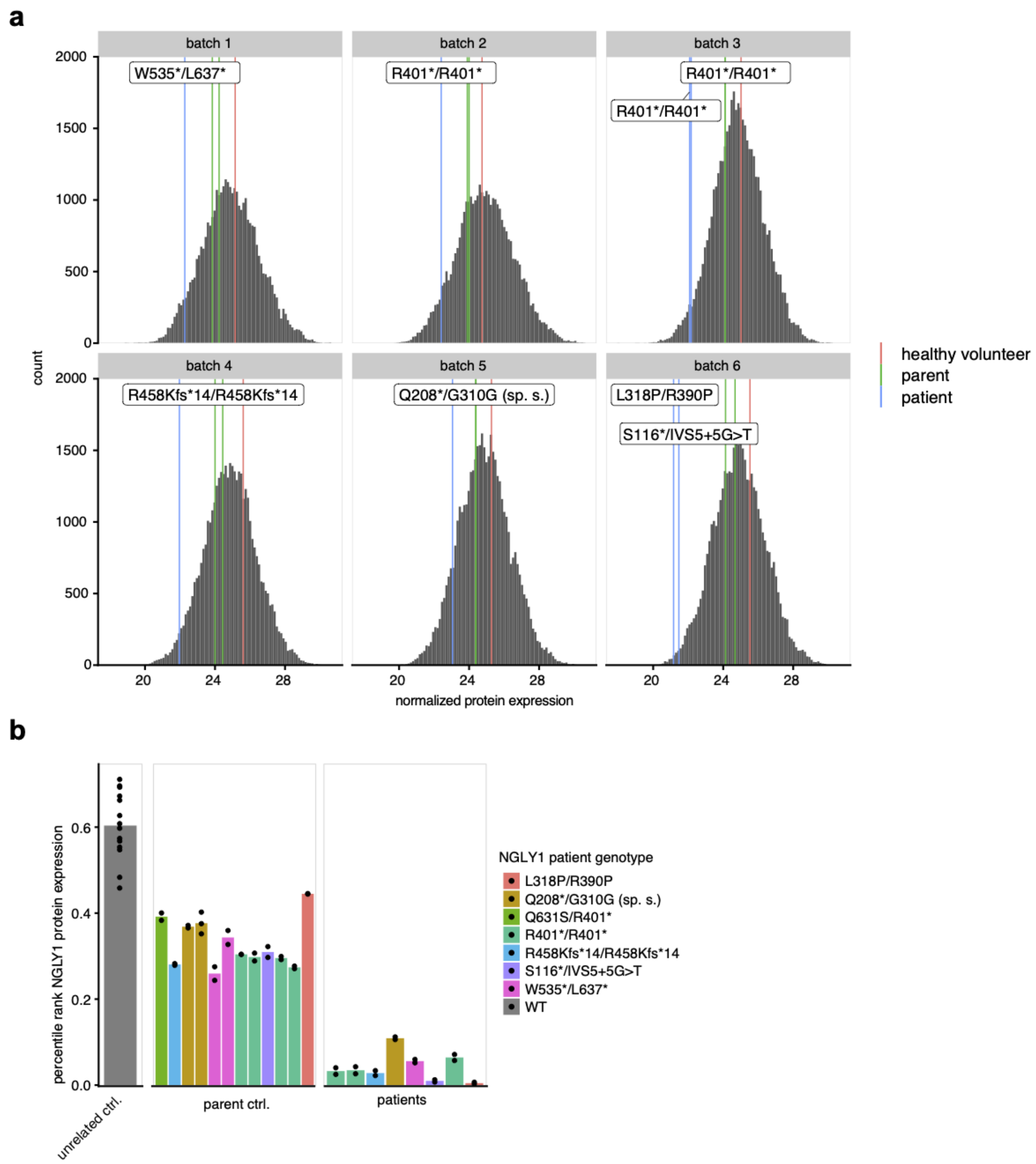


**Supplementary Figure 5. (a)** Histograms of normalized protein expression values for all detected proteins in LC-MS experiments in lymphoblastoid cell lines. Separate histograms are shown for each experimental batch. Vertical lines indicate NGLY1 protein expression values in patient (blue) and control cell lines (green are parent controls, red are unrelated controls). The vertical blue lines are labeled according to the corresponding patients' NGLY1 genotype. **(b)** Protein expression percentage rank of NGLY1 protein relative to all other detected proteins in lymphoblastoid cell lines. Dots represent independent samples and bars indicate sample means. Bars are colored according to the corresponding child patients' NGLY1 genotype.
